# Dissecting the roles of Haspin and VRK1 in Histone H3 threonine-3 phosphorylation during mitosis

**DOI:** 10.1101/2021.09.07.459242

**Authors:** Tyrell N. Cartwright, Rebecca J. Harris, Stephanie K. Meyer, Nikolaus A. Watson, Cheryl Tan, Fangwei Wang, Jonathan M.G. Higgins

## Abstract

Protein kinases that phosphorylate histones are ideally-placed to influence the behavior of chromosomes during cell division. Indeed, a number of conserved histone phosphorylation events occur prominently during mitosis and meiosis in most eukaryotes, including on histone H3 at threonine-3 (H3T3ph). At least two kinases, Haspin and VRK1 (NHK-1/ballchen in *Drosophila*), have been proposed to carry out this modification. Phosphorylation of H3 by Haspin has defined roles in mitosis, but the significance of VRK1 activity towards histones in dividing cells has been unclear. Here, using *in vitro* kinase assays, KiPIK screening, RNA interference, and CRISPR/Cas9 approaches, we were unable to substantiate a direct role for VRK1, or its homologue VRK2, in the phosphorylation of threonine-3 or serine-10 of Histone H3 in mitosis, although loss of VRK1 did slow cell proliferation. We conclude that the role of VRK1, and its more recently identified association with neuromuscular disease in humans, is unlikely to involve mitotic histone kinase activity. In contrast, Haspin is required to generate H3T3ph during mitosis.

## Introduction

The structure and function of chromosomes undergo radical changes during cell division, including chromosome condensation, downregulation of gene expression, removal of cohesion between chromosomes, and the assembly of kinetochores at centromeres to drive chromosome segregation. Protein kinases are instrumental in bringing about these alterations, and kinases that act directly on histones are well-placed to influence the behavior of chromatin. Indeed, a number of histone phosphorylation events occur prominently during mitosis and meiosis in most eukaryotes including yeasts, plants and animals (1). Among these are phosphorylation of histone H3 at threonine-3 (H3T3ph) and serine-10 (H3S10ph), and histone H2A at threonine-120 (H2AT120ph; or the equivalent H2AT119ph in *Drosophila*). For example, in mammalian cells, H3T3ph first becomes detectable at foci on chromosome arms in late G2/early prophase, concentrates most strongly at inner centromeres in prometaphase and metaphase, and declines during anaphase (2,3). A key function of H3T3ph is to position the Chromosomal Passenger Complex (CPC) correctly at inner centromeres (4-6). The CPC contains the kinase Aurora B that regulates chromosome segregation by modulating the phosphorylation status of a number of centromere and kinetochore substrates (7). In addition, H3T3ph may participate in a phospho-methyl switch to prevent histone reading proteins such as the transcription factor TFIID from binding to the adjacent modification H3K4me3 during mitosis (8).

Central to understanding the role of these histone phosphorylation events is knowledge of the kinases responsible. Two kinases have been proposed to carry out phosphorylation of H3T3 during mitosis. One of these is Haspin, an atypical kinase (encoded by Germ Cell-Specific Gene-2, GSG2) that phosphorylates H3 peptides, recombinant H3, and nucleosomes, specifically at threonine-3 *in vitro* (9,10). Haspin RNAi, inhibitors, or CRISPR/Cas9 targeting all remove mitotic H3T3ph in cultured mammalian cells, supporting the view that this is a critical kinase for H3T3ph during cell division (9,11-14). Loss or inhibition of Haspin impairs chromosome alignment and segregation in cell culture (9,12,15), particularly when cells are transiently blocked in mitosis (14), or when another kinase that helps recruit Aurora B to centromeres, Bub1, is also inhibited (16,17). In contrast, overall chromosome condensation is not noticeably altered by lowered Haspin activity (9).

Other work, however, has implicated Vaccinia-Related Kinase-1 (VRK1) as a mitotic H3T3 kinase. VRK1 is a member of a subgroup of the Casein Kinase-1 (CK1) family that also contains VRK2 and the pseudokinase VRK3 (18). VRK1 is reported to phosphorylate H3T3 *in vitro* (19-23), and VRK1 RNAi can diminish H3T3ph in mammalian cells (19,21,23,24). In some reports, but not others, VRK1 is also able to phosphorylate H3S10 *in vitro*, and to influence H3S10ph in cells (19-21,23-25). For example, infection of gastric epithelial cells with *Heliobacter pylori* was found to reduce H3T3ph and H3S10ph, and this could be rescued by overexpression of VRK1 (26). VRK1 overexpression causes nuclear condensation, leading to the suggestion that H3T3ph (and H3S10ph) are major regulators of chromosome condensation during mitosis (19), a model that has been widely disseminated (23,27-36).

The relative roles of Haspin and VRK1 in H3T3 phosphorylation remain to be defined. Some authors find that both VRK1 and Haspin are required for full H3T3ph in mitotic cells (21,24). Others have proposed that VRK1 is essential to trigger generalized H3T3ph in early mitosis, and that Haspin acts only later in mitosis. In this model, VRK1 phosphorylates H3T3ph to condense chromosomes and recruit the CPC, and Aurora B then activates Haspin to generate localized H3T3ph at centromeres (23,36). This work contrasts with other models in which the timing and location of H3T3ph is controlled indirectly by Cdk1, Bub1, Plk1 and Aurora kinases acting through Haspin and the H3T3ph phosphatase Repo-Man/PP1 (37-43). VRK1 mutations in humans appear to cause neurodegenerative disorders (30,44,45) and, consistent with this, VRK1^GT3/GT3^ hypomorphic mice have reduced brain weight and mild motor dysfunction (46). These findings reinforce the need to understand the molecular functions of VRK1. In this study, we wished to clarify the contribution of Haspin and VRK1 to H3T3ph in mitosis.

## Results

### Recombinant Haspin phosphorylates H3T3 more efficiently than VRK1 *in vitro*

To compare the activity of human Haspin and human VRK1 towards Histone H3T3, we conducted *in vitro* kinase assays using recombinant Haspin (residues 471-798), recombinant full-length VRK1, and peptides corresponding to the first 21 amino acids of H3. Three different preparations of recombinant VRK1, produced in both *E. coli* or in Sf9 insect cells, were similarly able to phosphorylate H3(1-21) at the high concentration of 10 nM. However, Haspin generated H3T3ph more strongly than VRK1, even at 0.1 nM (Figure 1A). VRK1 has been reported to phosphorylate nucleosomes more effectively than free histones (47), so we also compared the activity of Haspin and VRK1 on purified recombinant nucleosomes. In this case, Haspin but not VRK1 was able to phosphorylate H3T3 (Figure 1B). Therefore, in these *in vitro* assays, Haspin was at least 100-fold more active towards H3T3 than was VRK1.

**Figure 1.**
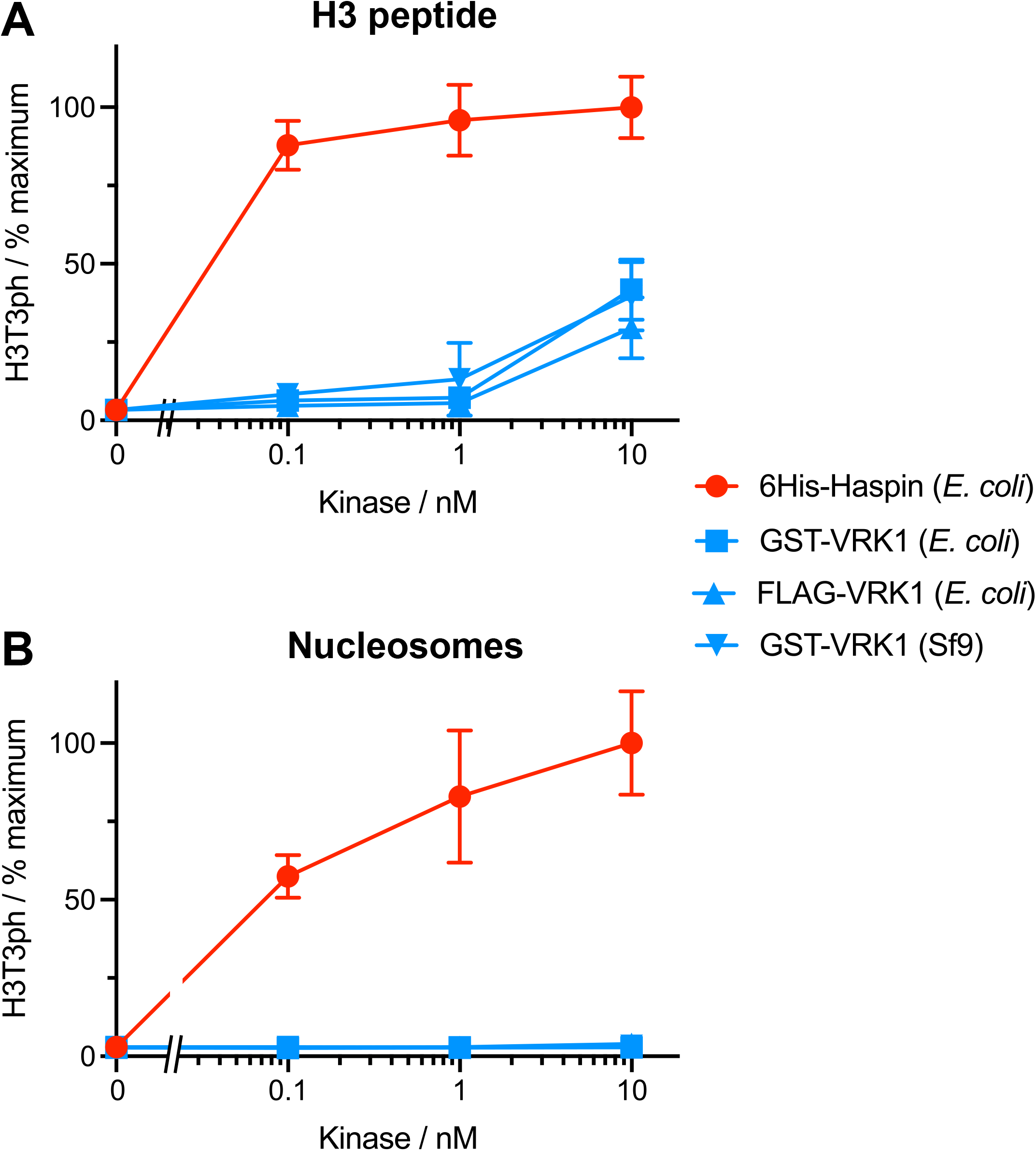
Phosphorylation of Histone H3 by recombinant human Haspin or VRK1 with **(A)** H3(1-21) peptide and **(B)** recombinant nucleosome substrates.

### A kinase activity screen implicates Haspin, not VRKs, as the mitotic H3T3 kinase

Recombinant kinases may lack cofactors (such as post-translational modifications or activating subunits) that are required for activity towards their substrates. To identify H3T3 kinases in a more physiological environment, and in an unbiased fashion, we made use of a new kinase screening methodology known as KiPIK (Kinase inhibitor Profiling to Identify Kinases) that uses cell extracts as a source of kinases (48). The method compares the pattern of inhibition of the unknown kinase activity in cell extracts with the inhibition profiles of known protein kinases determined *in vitro* using small molecule kinase inhibitors. We determined which kinase(s) in mitotic HeLa cell extract were able to phosphorylate H3(1-21) peptides, detected with anti-H3T3ph antibodies. To generate an inhibition fingerprint for H3T3 kinase activity in extract, we used a library of 140 kinase inhibitors that have been profiled on numerous purified kinases *in vitro* (Supplemental Table 1). This included 84 of the 178 inhibitors profiled at 0.5 µM on 300 kinases by Anastassiadis *et al*. (49), and 63 of the 158 inhibitors profiled at 1 µM or 10 µM on 234 kinases by Gao *et al*. (50), all using conventional *in vitro* kinase assays. In addition, it included all 72 inhibitors profiled on 388 different kinases by Davis *et al*. using the DiscoverX competition binding assay (51), and 50 of the 156 inhibitors tested at 10 µM for binding to 60 kinases by Federov *et al*. using a thermal denaturation assay (52). Together, this incorporates 444 kinases (405 unique human kinase genes; Supplemental Table 2).

The results showed that, in every case where Haspin was present in the profiling dataset, it was the top hit in the screen (Figure 2A-E, red circles). By contrast, no VRK kinase scored within the top 40% of kinases (Figure 2A-E, blue circles). Indeed, with the exception of the Anastassiadis screen (where VRK1 was 126^th^ out of 300 kinases), VRK1, VRK2 and VRK3 were all found in the bottom 20% of kinases (Figure 2A-E; Supplementary Table 3). These results are consistent with previous KiPIK screens for H3T3 kinases (48), where Haspin was the top overall hit, but VRK1 and VRK2 were never within the top 30% and were most often in the bottom 20% of kinases (Figure 2F; Figure S1). Of note, another kinase that has been proposed to phosphorylate a number of sites on H3 including H3T3, PASK (53), was also not a notable hit in these screens (Figure 2, Figure S1). Therefore, Haspin was reproducibly identified as the most likely mitotic H3T3 kinase in HeLa cell extracts.

**Figure 2.**
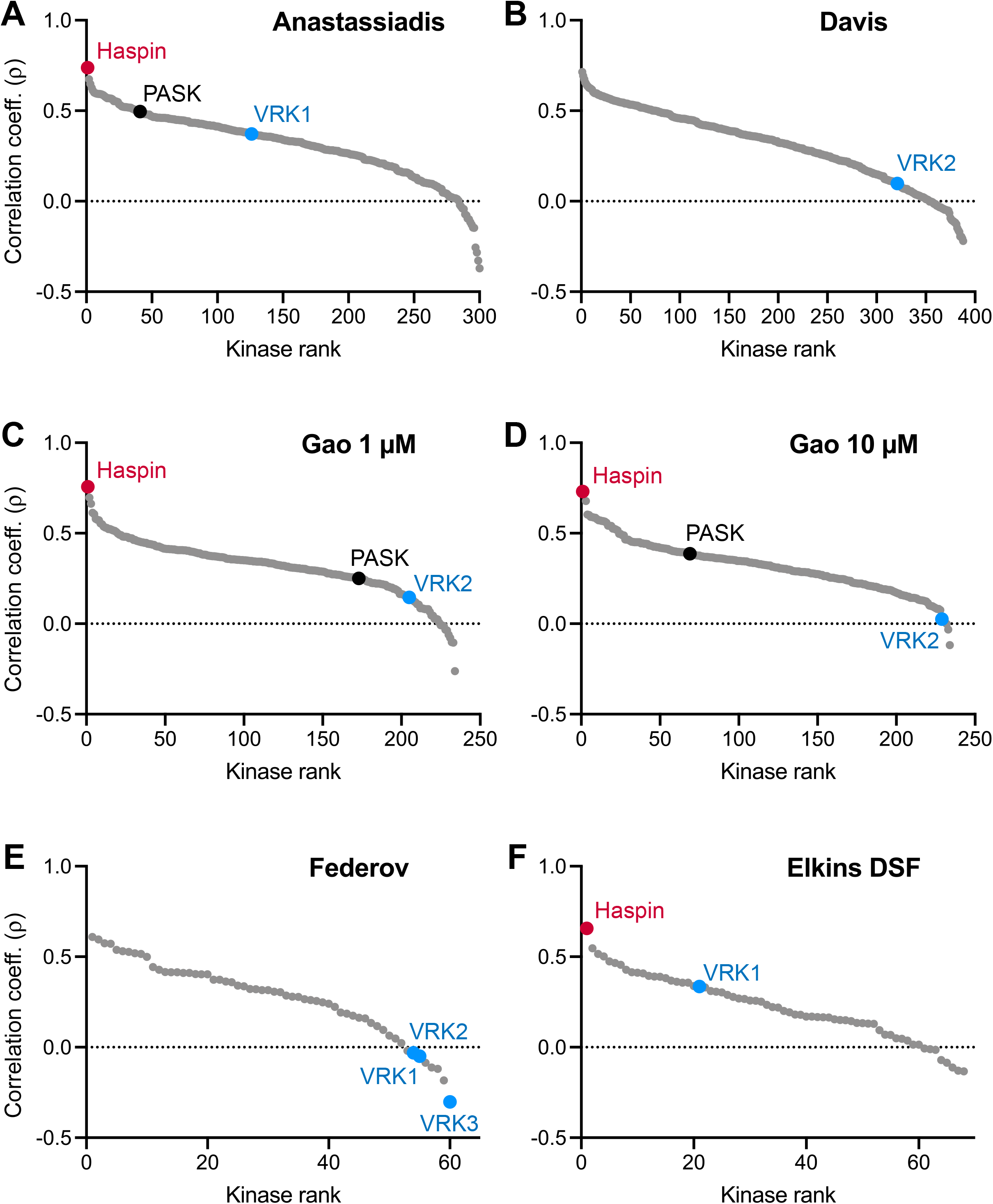
KiPIK screens for H3T3 kinase activity in mitotic HeLa cell lysates using the kinase profiling datasets of **(A)** Anastassiadis et al. (49), **(B)** Davis et al. (51), **(C, D)** Gao et al.,(50), **(E)** Federov et al. (52) and **(F)** the differential scanning fluorimetry (DSF) method of Elkins et al. (84). Some results of the Elkins DSF screen were previously published (48). The ranking of Haspin (red), VRK1, VRK2, VRK3 (blue) and PASK (black) are shown for all datasets in which they appear.

### A kinome-wide RNAi screen implicates Haspin, not VRKs, as the H3T3 kinase in mitosis

KiPIK screening uses cell extracts and appears to be relatively insensitive to the effects of upstream kinases in signaling cascades, and therefore is useful for identifying direct kinases for specific substrates (48). As an alternative unbiased approach to screen for H3T3 kinases in a cellular environment, and in manner that would allow the possible indirect effect of kinases to be observed, we analysed the results of a kinome-wide RNAi screen for H3T3 kinases in mitotic HeLa cells (48). Haspin was the top hit in this screen, and kinases known to regulate Haspin and the H3T3ph phosphatase Repo-Man-PP1 (i.e. Aurora B and Plk1 (37-39,41)) were also present in the top 5 kinase hits (Figure 3A). By contrast, none of the VRK kinases (nor PASK) were in the top 100 of the approximately 500 kinases tested. Therefore, RNAi screening also favours the hypothesis that Haspin, but not VRKs, are involved in the regulation of H3T3 in mitotic HeLa cells.

**Figure 3.**
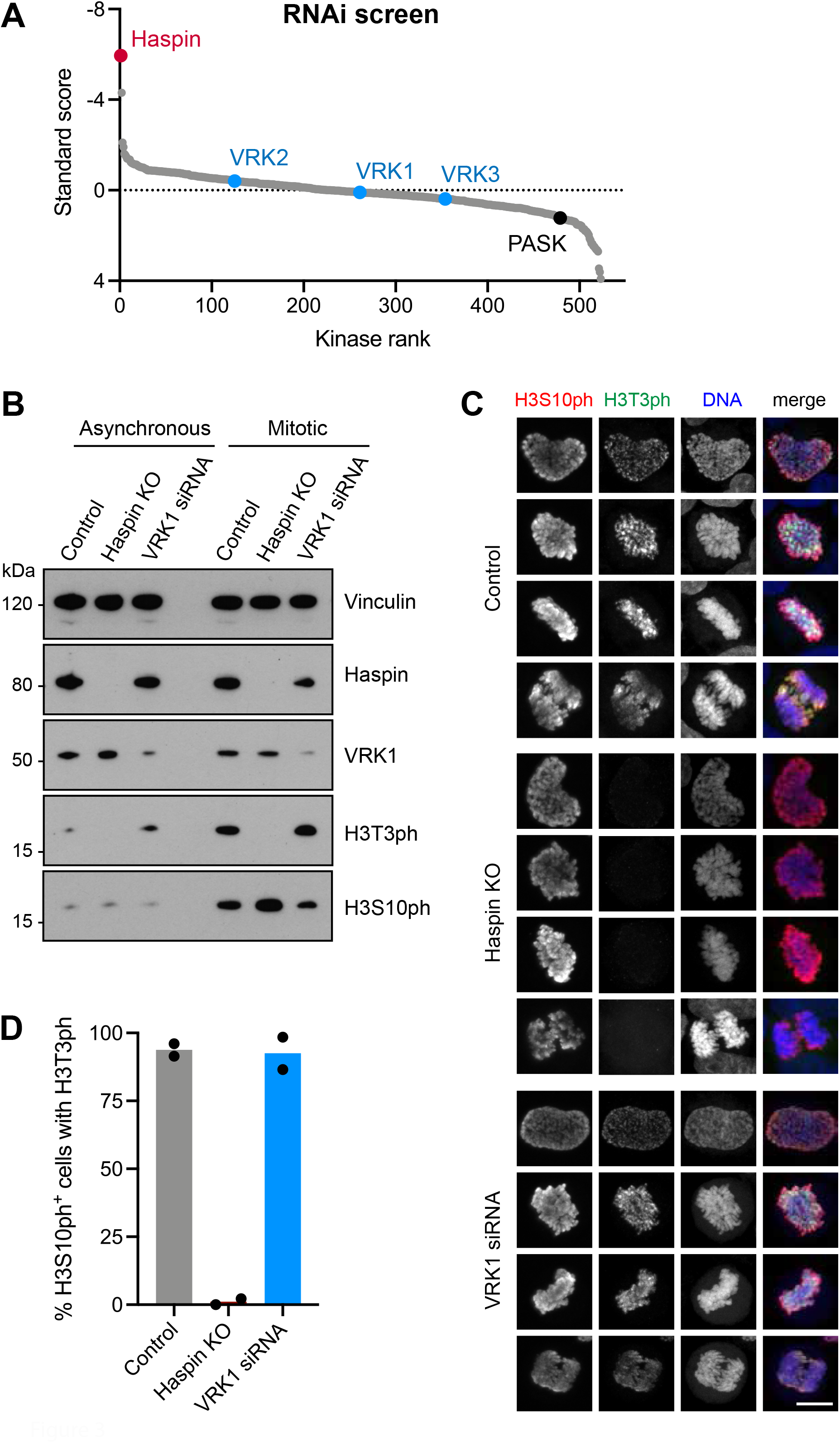
RNAi studies of VRK kinase function in mitosis. **(A)** Kinome-wide RNAi screen for kinases involved in H3T3 phosphorylation in HeLa cells. The standard score reflects the intensity of H3T3ph. Note that negative values indicate a decline in H3T3ph intensity. The ranking of Haspin (red), VRK1, VRK2, VRK3 (blue) and PASK (black) are shown. Some results of this screen were previously published (48). **(B)** RNAi depletion of VRK1 from HeLa cells does not cause a significant loss of H3T3ph as detected by immunoblotting. **(C)** RNAi depletion of VRK1 does not cause a significant loss of H3T3ph or H3S10ph in individual mitotic HeLa cells, as detected by immunofluorescence microscopy. Scale bar, 10 µm. **(D)** Quantification of the proportion of HeLa cells in (C) with H3S10ph (i.e. mitotic cells) that also stain for H3T3ph.

### Targeted RNAi for VRK1 does not reveal a role in H3T3 phosphorylation

Although the RNAi screen uses multiple pre-validated siRNAs against each kinase, we could not be certain that VRK1 was depleted in these experiments. We therefore conducted targeted experiments to deplete VRK1 from HeLa cells by RNAi. We were able to reduce the amount of VRK1 by approximately 80% +/-7% (n = 3). In immunoblotting experiments, this did not detectably reduce the extent of H3T3ph in asynchronous cells or in cells arrested in mitosis with the microtubule destabilizing inhibitor nocodazole (Figure 3B). Immunofluorescence microscopy also confirmed that VRK1 depletion did not detectably alter H3T3ph in mitosis (Figure 3C). By contrast, in HeLa cells lacking Haspin due to CRISPR/Cas9 targeting (14), H3T3ph could not be detected by immunoblotting or immunofluorescence microscopy (Figure 3B, 3C). As expected (9,12,13), loss of Haspin did not alter the extent of H3S10ph in HeLa cells detected by immunoblotting (Figure 3B) or immunofluorescence (Figure 3C). Consistent with this, loss of Haspin, but not VRK1, prevented detection of H3T3ph in mitotic cells containing H3S10ph (Figure 3D). VRK1 depletion caused a partial decrease in H3S10ph in immunoblotting experiments (Figure 3B), but not in individual mitotic cells as determined by immunofluorescence microscopy (Figure 3C). This is likely to be due to a small decrease in the proportion of VRK1 depleted cells blocked in mitosis by nocodazole compared to control cells (see below), although progression through mitosis itself was not obviously altered in asynchronously growing cells (Figure S2). Therefore, using RNAi in HeLa cells, we could not uncover a role for VRK1 in phosphorylation of H3T3, or H3S10.

### CRISPR/Cas9 targeting of VRK1 does not reveal a role in H3T3 phosphorylation

We were unable to remove all VRK1 from cells using RNAi, leaving open the possibility that low amounts of the kinase are able to maintain its putative role in H3 phosphorylation. To address this, we compared the effects of CRISPR/Cas9 targeting of Haspin and VRK1 in the haploid cell line HAP1 (see Figure S3). As expected, Haspin but not VRK1 could be detected in HAP1 VRK1 KO cells, and VRK1 but not Haspin could be seen in HAP1 Haspin KO cells (Figure 4A). As in HeLa cells, loss of Haspin eliminated H3T3ph detectable by immunoblotting of nocodazole-arrested mitotic HAP1 cells, but did not markedly affect H3S10ph (Figure 4B). By contrast, both H3T3ph and H3S10ph remained present in mitotic cells lacking VRK1 (Figure 4B). The partial decline in both H3T3ph and H3S10ph detected in mitotic cells by immunoblotting could be ascribed to the lower proportion of HAP1 VRK1 KO cells that accumulated in mitosis in nocodazole, as shown by the reduced intensity of Mitotic Protein Monoclonal-2 (MPM-2) phosphoepitopes seen by immunoblotting (Figure 4B), and by the lower mitotic index of nocodazole-treated HAP1 VRK1 KO cells compared to controls (Figure 4C). Indeed, live imaging and cell counting revealed that, although the duration of mitosis was only slightly increased in HAP1 VRK1 KO cells (Figure S4A), they did proceed more slowly through interphase than HAP1 control or Haspin KO cells (Figure 4D, Figure S4B), explaining why fewer VRK1 KO cells can accumulate in mitosis during exposure to nocodazole.

**Figure 4.**
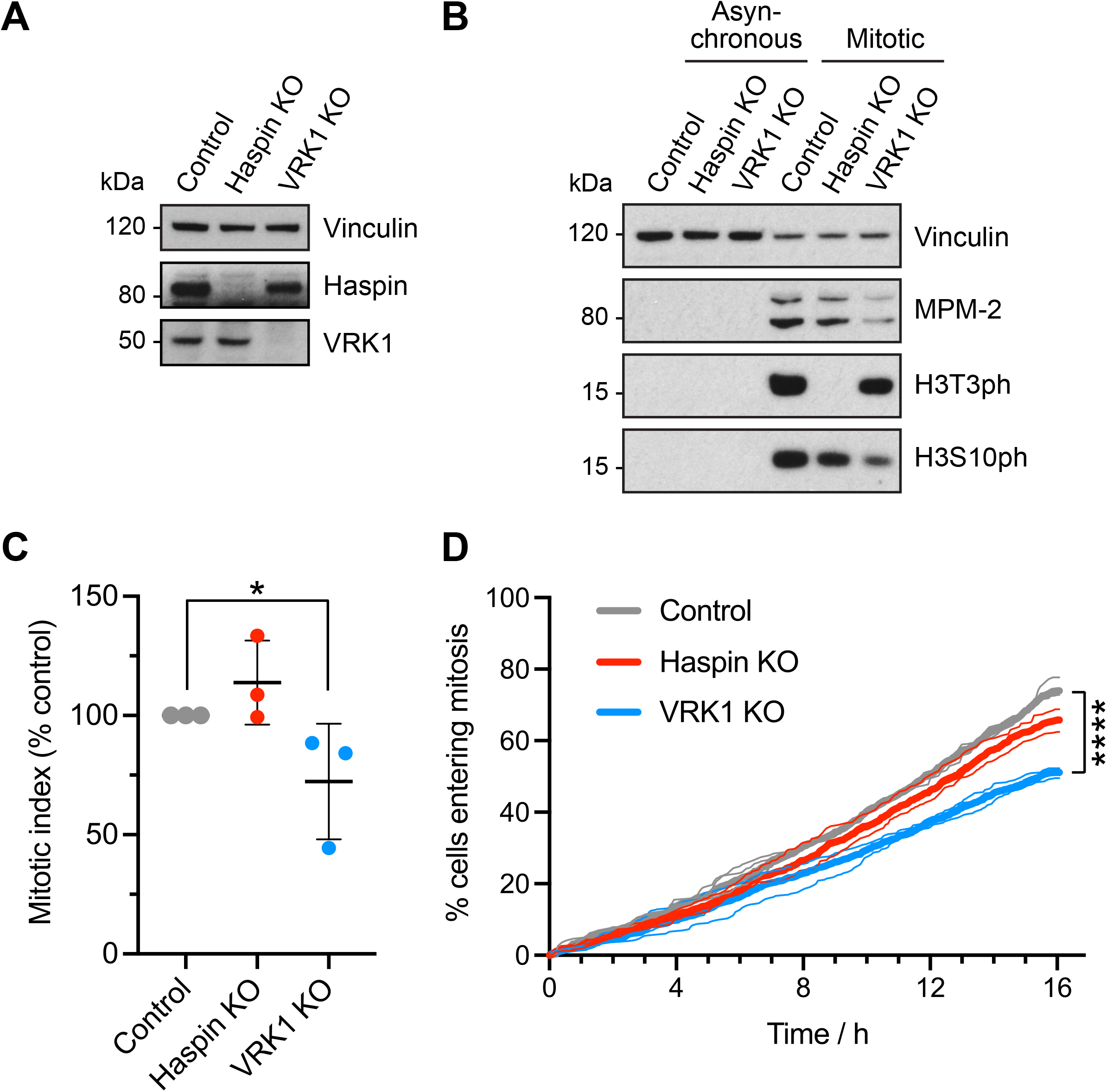
Histone H3 phosphorylation in control, Haspin KO and VRK1 KO HAP1 cells. **(A)** Immunoblotting shows the loss of Haspin in Haspin KO HAP1 cells, and the loss of VRK1 in HAP1 VRK1 KO cells. **(B)** Loss of VRK1 in HAP1 cells does not cause a significant loss of H3T3ph as detected by immunoblotting. **(C)** Control, Haspin KO and VRK1 KO HAP1 cells were allowed to accumulate in mitosis for 16 h in 300 nM nocodazole, and mitotic indices were then determined by immunofluorescence microscopy for MPM2. Error bars show means +/-SD. *, p = 0.045 (paired two-tailed Student t test using non-normalised data, n = 3). **(D)** Cumulative frequency of mitotic entry graphs for control, Haspin KO and VRK1 KO HAP1 cells observed by live imaging for 16 h. Between 193 and 555 cells were evaluated from three separate fields of view in each of three independent experiments, shown as thin lines. The combined results are shown as thick lines. Statistical significance was determined from the combined data using Kaplan-Meier curve analysis and a Mantel-Cox log rank test (****, p < 0.0001). Mean cell cycle duration for Control, 24 h; Haspin KO, 26 h; and VRK1 KO, 33 h; 95% confidence intervals [23.5, 24.0], [26.0, 26.6] and [32.5, 33.0] respectively, as determined by linear regression.

This interpretation was supported by quantifying the intensity of H3 phosphorylation in individual mitotic cells by immunofluorescence microscopy. Haspin KO HAP1 cells lost H3T3ph, but the intensity of H3T3ph in HAP1 VRK1 KO cells appeared unchanged, and H3S10ph appeared unaffected in both cell lines (Figure 5). Quantification of the number of cells with H3S10ph that also had H3T3ph confirmed this result (Figure S5A). These experiments also supported the view that progression through mitosis was not substantially altered by loss of Haspin or VRK1 (Figure S5B). These results reveal that eliminating detectable VRK1 from HAP1 cells does not prevent normal H3T3 (or HS10ph) phosphorylation in mitosis.

**Figure 5.**
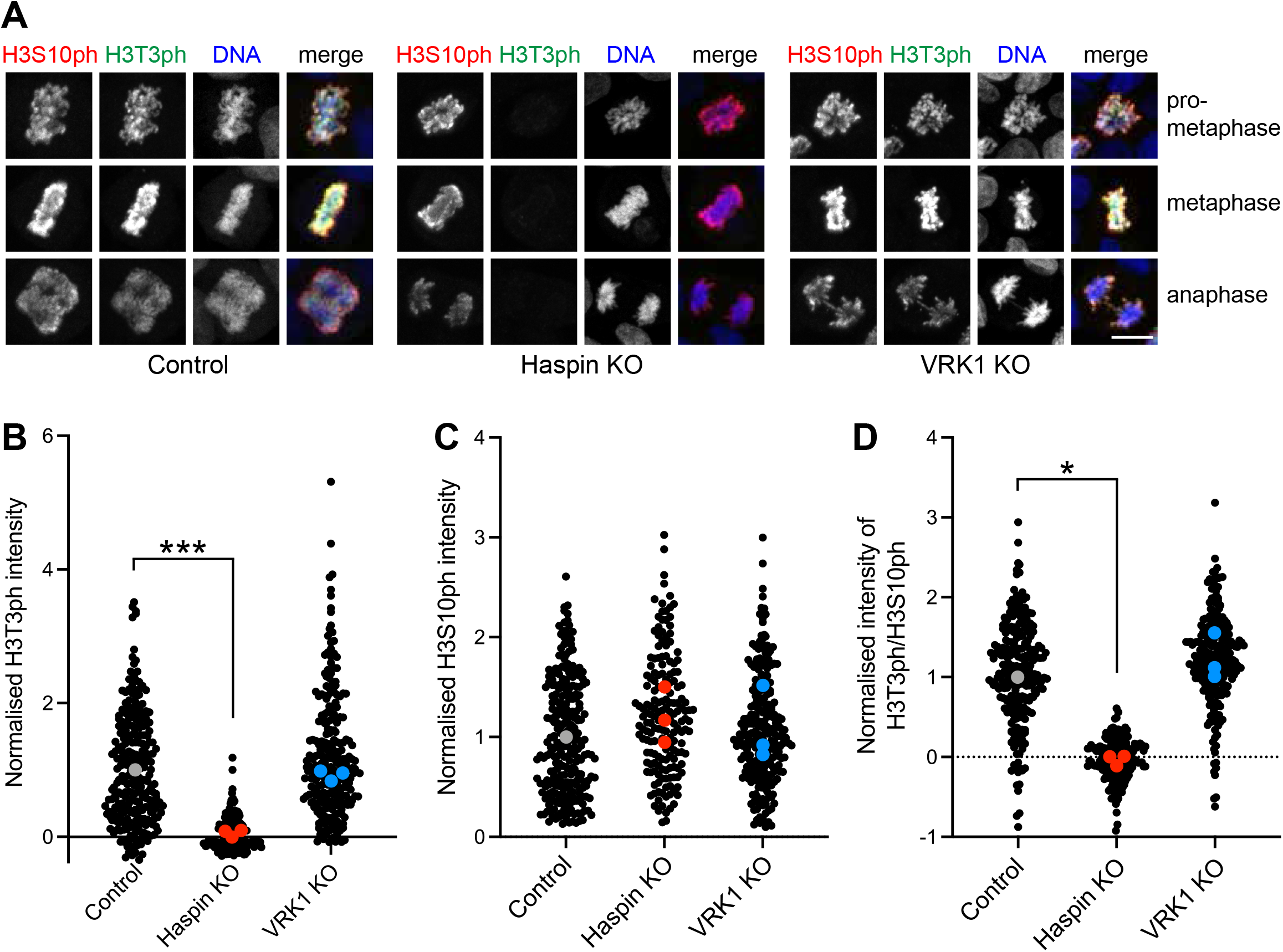
**(A)** Loss of Haspin, but not VRK1, eliminates H3T3ph in individual mitotic HAP1 cells, as detected by immunofluorescence microscopy. H3S10ph appears unaffected in both cases. Scale bar, 10 µm. **(B)** Quantification of the intensity of H3T3ph staining in cells as in (A). Black symbols represent individual cells. Grey, red and blue symbols show the average values for each of 3 independent experiments, normalised to control HAP1 cells. ***, p = 0.0017 (two-tailed t test using non-normalised data; n = 3). **(C)** Quantification of the intensity of H3S10ph staining in cells as in (B). Two-tailed t tests using non-normalised data revealed no significant differences at p = 0.05 (n = 3). **(D)** Quantification of the ratio of H3T3ph intensity to H3S10ph intensity in cells as in (B). *, p = 0.0499 (two-tailed t test using non-normalised data; n = 3).

### RNAi does not reveal a role for VRK2 in H3T3 phosphorylation

Human cells contain a second VRK1-related protein with kinase activity, VRK2. To determine if H3T3ph remaining in HAP1 VRK1 KO cells could be due to the presence of VRK2, we depleted VRK2 from these cells by RNAi. We were able to reduce VRK2 protein expression levels by 84% +/-10% (n = 3). In immunoblotting experiments, this did not detectably reduce the extent of H3T3ph or H3S10ph in cells arrested in mitosis with the microtubule destabilizing inhibitor nocodazole (Figure 6), suggesting that VRK2 is not responsible for the persistence of H3T3ph in HAP1 VRK1 KO cells. As discussed earlier, HAP1 VRK1 KO cells had slightly reduced levels of H3T3ph, H3S10ph, and MPM2 phosphoepitopes, most likely explained by the decline in the proportion of mitotic cells.

**Figure 6.**
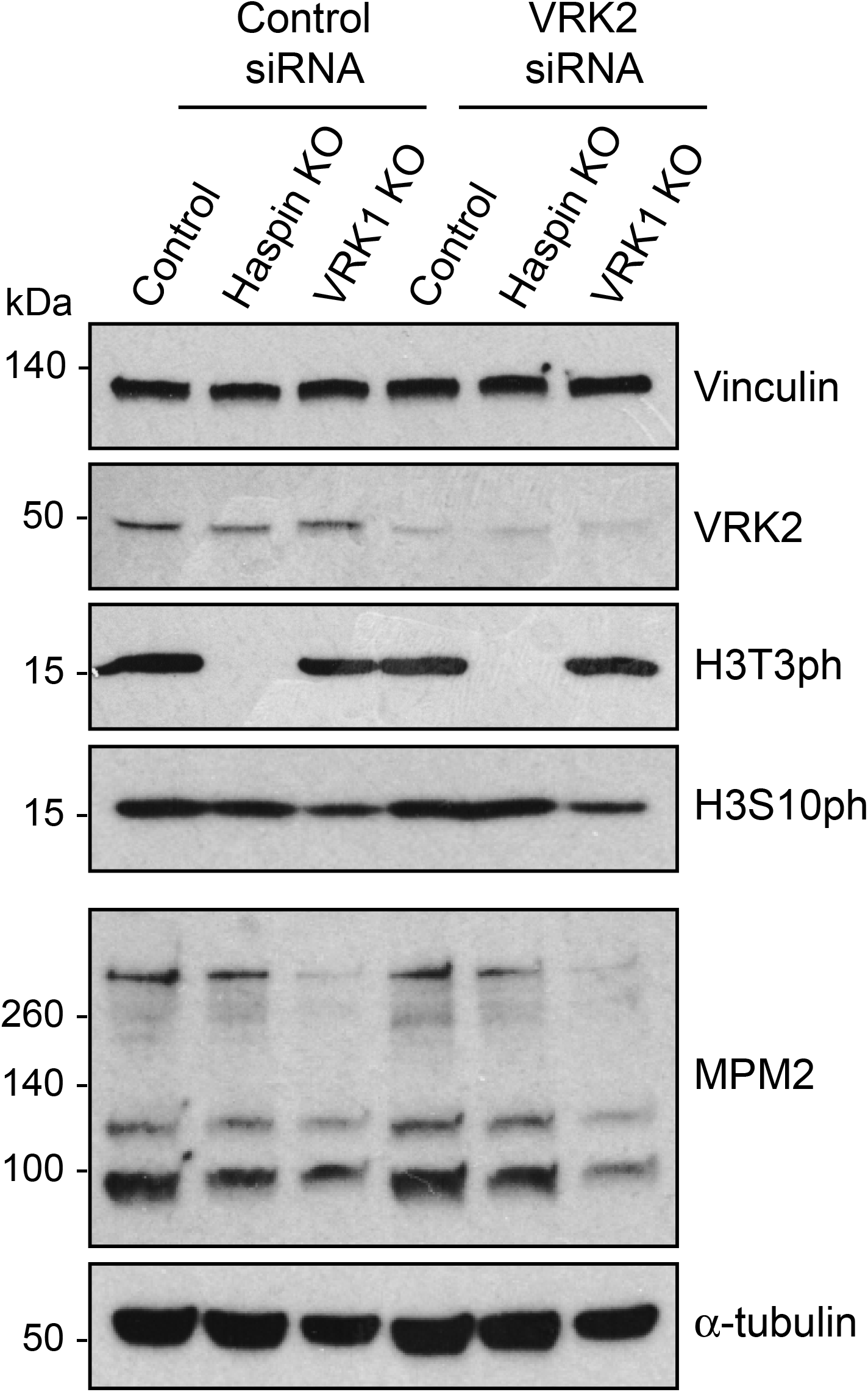
RNAi of VRK2 does not have a detectable influence on H3T3ph. Control and VRK1 KO HAP1 cells were treated with control or VRK2 siRNAs and subjected to immunoblotting with the stated antibodies. Note that the top 4 panels are from an immunoblot different from the one in the bottom 2 panels.

## Discussion

Here, we find that Haspin is a much more potent kinase than VRK1 for H3T3 *in vitro*. In addition, using KiPIK assays in mitotic HeLa cell extracts, none of the VRK family kinases are found to be strong candidates for the H3T3 kinase in mitosis while, in contrast, Haspin is the highest ranked kinase. Similarly, in a kinome-wide RNAi screen for mitotic H3T3ph kinases, Haspin was the top hit, while none of the VRKs were in the top 100. CRISPR/Cas9 targeting of Haspin, but not VRK1, causes loss of detectable H3T3ph during mitosis in HAP1 cells. Furthermore, depleting VRK2 in combination with loss of VRK1 does not lead to detectable reductions in H3T3ph, suggesting that redundancy with VRK2 does not underlie our failure to influence H3T3ph when VRK1 expression is ablated. VRK3 is unlikely to contribute as it is a pseudokinase with a degraded active site structure (31) that is unable to bind divalent cations or ATP (54), and is unable to phosphorylate casein or H3 (20,55). In addition, PASK did not score highly as a mitotic H3T3 kinase in either KiPIK or RNAi screens, concordant with the finding that CRISPR/Cas9 targeting of the PASK gene did not markedly alter H3T3ph in mouse C2C12 cells (53).

These results suggest that Haspin is the major H3T3 kinase during mitosis in human cells. This is consistent with reports that Haspin is required for normal mitotic H3T3ph in mouse cells (3), sea urchin embryos (56), fruit flies (57,58), *Arabidopsis* (59,60), and fission yeast (6), and for H3T3ph during meiosis in mouse oocytes (61,62) and frog egg extracts (5). Indeed, Haspin orthologues are present in all organisms that are reported to have H3T3ph in mitosis to date. By contrast, VRK family kinases are present in many metazoa (eg VRK-1 in *C. elegans*, and NHK-1/ballchen in *Drosophila*) but appear absent from plants and yeasts (28,55).

It remains uncertain why previous experiments have suggested a role for VRK1 in producing H3T3ph in mitosis. However, multiple reports indicate that VRK1 influences cell cycle progression. VRK1 regulates the expression of Cyclin D1, and VRK1 depletion impedes progression from G1 to S phase in cultured human cells (35,63-65). This could hinder cells reaching mitosis and so lead to a loss of modifications that are abundant during this time, such as H3T3ph (and H3S10ph). Indeed, although it is clear that HAP1 cells lacking detectable VRK1 can proliferate in culture, live imaging revealed that a lower proportion of VRK1 KO cells enter mitosis per unit time, consistent with a delay in the cell cycle outside mitosis, such as during G1. In addition, a previous report found that a high proportion of HeLa cells die in mitosis following VRK1 RNAi (66). The loss of H3 phosphorylation seen in previous immunoblotting experiments that used nocodazole to accumulate mitotic cells, therefore, may be an indirect effect of a reduced mitotic index. By contrast, in experiments where the intensity of H3T3ph (and H3S10ph) in individual mitotic cells was measured, we found no evidence for a decline in these phosphorylation marks.

Analysis of VRK1 function in cell division in model organisms is complicated because complete loss of function is lethal in mice, flies and worms (34,67,68), so only hypomorphic alleles or the results of partial depletions have been examined. Male and female VRK1^GT3/GT3^ gene-trapped mice are sterile, indicating defects in gametogenesis. Indeed, there is a strong defect in proliferation of spermatogonial cells in male mice, commensurate with their reduced levels of cell cycle markers such as Proliferating Cell Nuclear Antigen (PCNA). The reduction in H3S10ph in these testes is most likely due to the reduced numbers of proliferating cells, not a specific loss of H3S10ph due to reduced VRK1 function (32). Consistent with this, sterility of female VRK1^GT3/GT3^ mice appears to be due to meiotic defects in oocytes, but these oocytes have normal levels of H3S10ph (69).

Hypomorphic NHK-1 mutant flies are also female-sterile, and have defects in clustering and condensation of chromosomes in oocytes (67,70). Mitotic chromosome condensation and segregation are abnormal in NHK-1-depleted *Drosophila* S2 cells and in NHK-1 mutant fly larvae, as well as in VRK-1-depleted *C. elegans* embryos (67,68,71). Notably, increases in the number of mitotic cells are common in hypomorphic NHK-1 mutant fly embryos and larvae, consistent with prolonged mitosis. In contrast, the proportion of mitotic cells *decreases* in embryos with lethal NHK-1 mutations (67,70,71), suggesting cell cycle defects prior to mitosis. However, in both cases, H3S10ph levels appear normal in individual mitotic cells, arguing against a direct role for NHK-1 in phosphorylating H3S10 (71). Similar results were obtained in cultured human cells arrested in mitosis, where VRK1 RNAi did not influence H3S10ph (72).

Putting together these findings, the reason that alterations in VRK1 activity give rise to different cell cycle defects in different systems may be due to the different mutants analysed and varying degrees of depletion obtained in different experiments. In addition, the extent to which other kinases such as VRK2 can act redundantly may vary depending on the organism, tissue or function in question. Indeed, there is a second VRK-related kinase in *Drosophila*, CG8878, although it has a distinctive split kinase domain and its molecular function remains unclear (73). Nevertheless, there is strong evidence that VRK1 deficiency affects both mitosis and meiosis. However, our results suggest these effects are unlikely to be due to changes in H3T3ph phosphorylation. In addition, H3S10ph does not appear to be directly affected by loss of VRK1, in line with other studies discussed above (32,71,72). Interestingly, fly NHK-1 and human VRK1 have also been reported to phosphorylate nucleosomal H2AT120 (H2AT119 in flies) (47,65). Phosphorylation of this site is clearly elevated in mitosis and meiosis in human cells, flies, budding and fission yeast (47,70,74,75). A female-sterile mutation of NHK-1 diminishes H2AT119ph in fly oocytes (70), but whether this is a direct or indirect effect of NHK-1 on H2A phosphorylation remains unknown. Ooctyes of VRK1^GT3/GT3^ mice have normal levels of H2AT120ph (69). Moreover, neither mutation nor depletion of NHK-1 prevents mitotic H2AT119ph in flies or *Drosophila* S2 cells (70,74). Instead, the kinetochore-binding kinase Bub1 is clearly vital for H2AT120ph in budding and fission yeast, mice, and human cells (75-77). Together, there is little evidence to support a significant role for VRK1 in direct phosphorylation of H3T3ph or other histone residues in mitosis.

VRK1 phosphorylates and inhibits Barrier-to-Autointegration Factor (BAF) to regulate nuclear envelope dissolution during mitosis and meiosis in flies, worms and mammalian cells (66,68,71,72,78,79), and this seems likely be a major contributor to its cell division phenotypes. In addition, we have not ruled out a role for VRK1 in histone phosphorylation outside mitosis. For example, worm VRK-1 is expressed in post-mitotic cells and can influence lifespan by activating AMP-activated protein kinase (80), and Aihara et al. have suggested that VRK1 regulates gene expression by phosphorylating H2AT120 at gene promoters such as *CCND1*/Cyclin D1 in mice (65). A number of H3S10 kinases that act to regulate transcription in interphase cells have been characterized, including MSK1/2 (28,33). We are not aware of H3T3 kinases that act during interphase in vertebrates, but Haspin itself may have interphase roles in *Drosophila* (58), and the plant-specific MUT9 kinases generate H3T3ph in *Chlamydomonas* and *Arabidopsis* and appear to be involved in epigenetic silencing (81,82). Interestingly, like the VRK family, MUT9 kinases are part of the CK1 superfamily and it is conceivable that VRKs serve a similar function in metazoa. It also will be interesting in future to determine if the mitotic or non-mitotic functions of VRK1 underlie the neuromuscular disorders observed in mice and human patients (30,44-46).

## Methods

### Cell culture and transfection

HeLa cells, HeLa clone D2 Haspin KO cells (14), and parental controls, were maintained in DMEM with 10% (v/v) FBS, 100 U/ml penicillin and streptomycin and 2 mM L-glutamine at 37°C and 5% CO_2_ in a humidified incubator. HAP1 knock out cell lines for GSG2 (Haspin; HZGHC000047c016) and VRK1 (HZGHC000073c014), as well as a parental control line (C631), were obtained from Horizon Discovery (Cambridge, UK) and maintained in IMDM with 10% (v/v) FBS, 100 U/ml penicillin and streptomycin and 2 mM L-glutamine at 37°C and 5% CO_2_ in a humidified incubator. HeLa D2 parental control cells were transfected with 300 ng/ml human VRK1 MISSION® esiRNA (EHU119541, Sigma) using HiPerFect Transfection Reagent (301707, Qiagen) and analysed after 24 h. HAP1 cells were transfected with 600 ng/ml human VRK2 MISSION® esiRNA (EHU070731, Sigma-Aldrich) using Lipofectamine RNAiMAX (13778075, ThermoFisher) and analysed after 24 h. MISSION® siRNA Universal Negative Control (SIC001, Sigma-Aldrich) was used at an equal concentration as the control for RNAi experiments. Where stated, cells were blocked in mitosis by treatment with 300 nM nocodazole (Sigma) for 16 h.

### Cell counting

HAP1 cell lines were counted using trypan blue (Sigma) and a haemocytometer, seeded at 100,000 cells/well in 6-well plates and recounted at 24 h intervals.

### SDS-PAGE and immunoblotting

Cells were washed once in PBS and then lysed in 141 mM Tris Base, 106 mM Tris HCl, 2% LDS, 1% Benzonase nuclease (E1014, Millipore), pH 8.5, at room temperature for 15 min and snap frozen in liquid nitrogen. The protein concentration was determined using Rapid Gold BCA Protein Assay Kit (Pierce). Cell lysates (20 µg) were run on NuPAGE 4-12% Bis-Tris protein gels (Invitrogen) using standard procedures, transferred to 0.2 µm polyvinylidene fluoride membranes and blocked in 5% milk in TBST (0.05% Tween 20 in Tris-buffered saline) for 1 h at room temperature before overnight incubation with primary antibodies. Membranes were washed thrice in TBST then incubated with a HRP conjugated secondary antibody for 1 h at room temperature in TBST with 5% milk, and signals were detected by chemiluminescence (Clarity ECL, Biorad). Antibodies used in immunoblotting were as follows: rabbit polyclonal anti-Vinculin (Cell Signaling Technology, #13901), anti-Haspin (Abcam, ab21686, lot 156753), and anti-H3T3ph (B8634; (9)) with anti-rabbit IgG HRP (Cell Signaling Technology, #7074) and mouse monoclonal anti-VRK1 (Abcam, ab171933), anti-VRK2 (Santa Cruz, sc-365199), anti-H3S10ph (Millipore, 05-806), anti-α-tubulin (B-5-1-2, Sigma, T5168) and anti-MPM2 (Millipore, 05-368) with anti-mouse IgG HRP (Cell Signaling Technology, #7076).

### Immunofluorescence

Cells were fixed for 10 min with 4% paraformaldehyde in PBS, washed twice in PBS, then permeabilized for 5 min with 0.5% Triton X-100 in PBS. After washing twice in PBS, cells were incubated for 1 h in 5% BSA in PBS, 0.05% Tween20 at room temperature, and then with primary rabbit antibody against H3T3ph (B8634) and, in some cases, with mouse anti-MPM2, at 37°C in 5% BSA in PBS, 0.5% Tween20. After washing twice with 0.05% Tween20 in PBS, cells were incubated for 1 h with fluorophore-conjugated secondary antibody anti-rabbit-IgG Alexa Fluor®488 (Invitrogen A32731) and, where necessary, anti-mouse IgG-Alexa Fluor®594 (Invitrogen, A-32744) at 37°C in 5% BSA in PBS, 0.5% Tween20. After washing twice with PBS, 0.5% Tween20, cells were incubated with rabbit phospho-Histone H3 (Ser10) (D2C8) XP® mAb directly conjugated to Alexa Fluor®647 (Cell Signaling Technology, #3458) for 1 h at room temperature. After washing twice with PBS, 0.5% Tween20, and once with milliQ H_2_O, samples were mounted using ProLong™ Diamond Antifade Mountant with DAPI (ThermoFisher, P36962). Images were captured on a Nikon A1 confocal microscope with a 20x Air/NA 0.75 or 60x Oil/NA 1.4 objectives using Nikon Elements software.

### Quantification of immunofluorescence

To quantify immunofluorescence intensities in individual cells, and to determine mitotic indices, images were analysed in Fiji 2.0 (83). Cells were identified from DAPI staining by automatic thresholding of maximum intensity projections and binarization followed by use of the watershed tool to separate adjacent cells. Cells were then counted using the analyse particles tool with a minimum cut off of 20 µm^2^. The fluorescence intensities for H3T3ph and H3S10ph in each cell was quantified using sum intensity projections. Mitotic cells were identified as those whose fluorescence intensity for H3S10ph classified them as outliers using the iterative Grubbs’ method (α = 0.0001) in Prism 9.0.2 (GraphPad). Visual inspection of DAPI stained cells confirmed that, for H3S10ph, this included essentially all mitotic cells. Automatically identified particles containing more than one mitotic cell were eliminated from the analysis. Background fluorescence was determined from the average fluorescence intensity of all non-mitotic cells. MPM2 positive cells were identified similarly with the additional use of the fill holes tool prior to watershedding.

### Live imaging

Live imaging was performed in glass-bottomed FluoroDishes (WPI) in IMDM medium (12440053, ThermoFisher). DNA was stained with 25 nM SiR-DNA (Spirochrome). DIC and SiR-DNA images at 6 z-steps per field (2 µm step size) were acquired at 4 min intervals on a Nikon A1R confocal microscope equipped with a 20x 0.75 NA PlanApo VC DIC objective using Elements v5.22 software (Nikon, Japan). In a blinded fashion, the time of mitotic entry was determined as the earliest time point at which the initiation of cell rounding was observed by DIC and/or the time point prior to nuclear envelope breakdown observed by SiR-DNA staining, and the time of mitotic exit was defined as the first frame in which anaphase was apparent.

### KIPIK screens

To prepare mitotic cell extract, HeLa cells were treated with 300 nM nocodazole for 15 h, collected by shake off, and lysed at 4°C in 50 mM Tris, 0.25 M NaCl, 0.1% Triton X100, 10 mM MgCl_2_, 2 mM EDTA, 1 mM DTT, pH 7.5 with protease inhibitor cocktail (Sigma P8340), PhosSTOP (Merck), 1 mM PMSF, 0.1 µM okadaic acid, 10 mM NaF, 20 mM β-glycerophosphate, at approximately 30 × 10^6^ cells/ml. Extracts were immediately flash frozen in liquid nitrogen. Kinase reactions contained 10 µM kinase inhibitor or DMSO (vehicle control), 0.35 µM Histone H3(1-21) peptide (ARTKQTARKSTGGKAPRKQLA-GGK-biotin; Abgent), 0.2 mM ATP and 5% cell extract, in KiPIK buffer (50 mM Tris, 10 mM MgCl_2_, 1 mM EGTA, 10 mM NaF, 20 mM β-glycerophosphate, 1 mM PMSF, pH 7.5 with PhosSTOP (Merck)). Reactions were carried out in duplicate in 384-well microplates (ThermoFisher), at 37°C for 30 min, in a total volume of 35 µl/well. An expanded custom library of inhibitors similar to that described previously (48) was used (Supplementary Table 1). For detection of phosphorylation, High Capacity Streptavidin-coated 384-well plates (Pierce) were washed thrice with TBS, 0.1% Tween20, then, 35 µl/well of completed KiPIK extract kinase reaction was added and incubated at room temperature for 1 h. After washing, 0.2 µg/ml H3T3ph B8634 antibodies were added at 40 µl/well in TBS, 0.1% Tween20, 5% BSA, for 1 h at room temperature. After washing, 40 µl/well of HRP-conjugated anti-rabbit IgG antibodies (Cell Signaling Technology, #7074) in TBS, 0.1% Tween20, 5% BSA were added for 1 h at room temperature. After washing, HRP-conjugated antibody binding was detected using TMB substrate (New England Biolabs) according to the manufacturer’s instructions. Pipetting was performed with a Biomek FX liquid handling robot (Beckman Coulter). Absorbance readings were made using a Polarstar Omega microplate reader (BMG Labtech).

### KIPIK results analysis

Using the mean of duplicate determinations, a standard score reflecting kinase activity in the presence of each inhibitor was calculated where standard score = (mean absorbance in presence of inhibitor - mean absorbance of multiple DMSO controls)/standard deviation of DMSO controls. The inhibition by each compound was defined as: %inhibition = 100 × (standard score for inhibitor/lowest standard score on plate). The %inhibition scores for all inhibitors were then compiled to produce the inhibition fingerprint of H3T3 phosphorylation. Using Excel (Microsoft), Pearson’s correlation (ρ) was then calculated for this fingerprint against the inhibition profiles of each of the kinases profiled *in vitro* against that inhibitor library, excluding mutant kinases. We calculated z-scores where z = (observed ρ – mean ρ of null distribution) / standard deviation of null distribution as described (48).

### *In vitro* kinase assays

*In vitro* kinase reactions contained the stated kinase concentrations, 0.35 µM H3(1-21)-GGK-biotin peptide (Abgent) or 0.35 µM recombinant biotinylated human mononucleosomes (16-0006, Epicypher) and 0.2 mM ATP in KiPIK buffer. Reactions were carried out in triplicate in 384-well microplates (ThermoFisher), at 37°C for 30 min, in a total volume of 35 µl/well. The kinases used were: full-length GST-VRK1 produced in *E. coli* (P5776, Abnova), FLAG-VRK1 produced in *E. coli* (31243, Active Motif), GST-VRK1 produced in insect Sf9 cells (ab125555, Abcam) and 6His-Haspin residues 471-798 produced in *E. coli* (9). Phosphorylation was detected as described for KiPIK experiments above.

### Statistical Analysis

Statistical analyses were carried out using Prism 9.0.2 (Graphpad).

## Contributions

TNC and RH designed, conducted and analysed multiple experiments; SKM carried out KiPIK screens; NAW conducted KiPIK and RNAi screens; CT conducted RNAi experiments; FW provided Haspin KO HeLa cells; JMGH supervised the project, analysed data, and wrote the manuscript draft. All authors made comments on the manuscript.

## Acknowledgements

The PKIS1 library was supplied by GlaxoSmithKline LLC and the Structural Genomics Consortium under an open access Material Transfer and Trust Agreement: http://www.sgc-unc.org. This study was funded by a Wellcome Trust Investigator Award (106951/Z/15/Z), a Royal Society Wolfson Research Merit Award, and an MRC Confidence in Concept award to JMGH, and a Royal Society Newton Advanced Fellowship (NA140075) to FW and JMGH. For the purpose of open access, the author has applied a CC BY public copyright licence to any Author Accepted Manuscript version arising from this submission.

