## Supplemental Figures for "Dissecting the roles of Haspin and VRK1 in Histone H3 threonine-3 phosphorylation during mitosis"

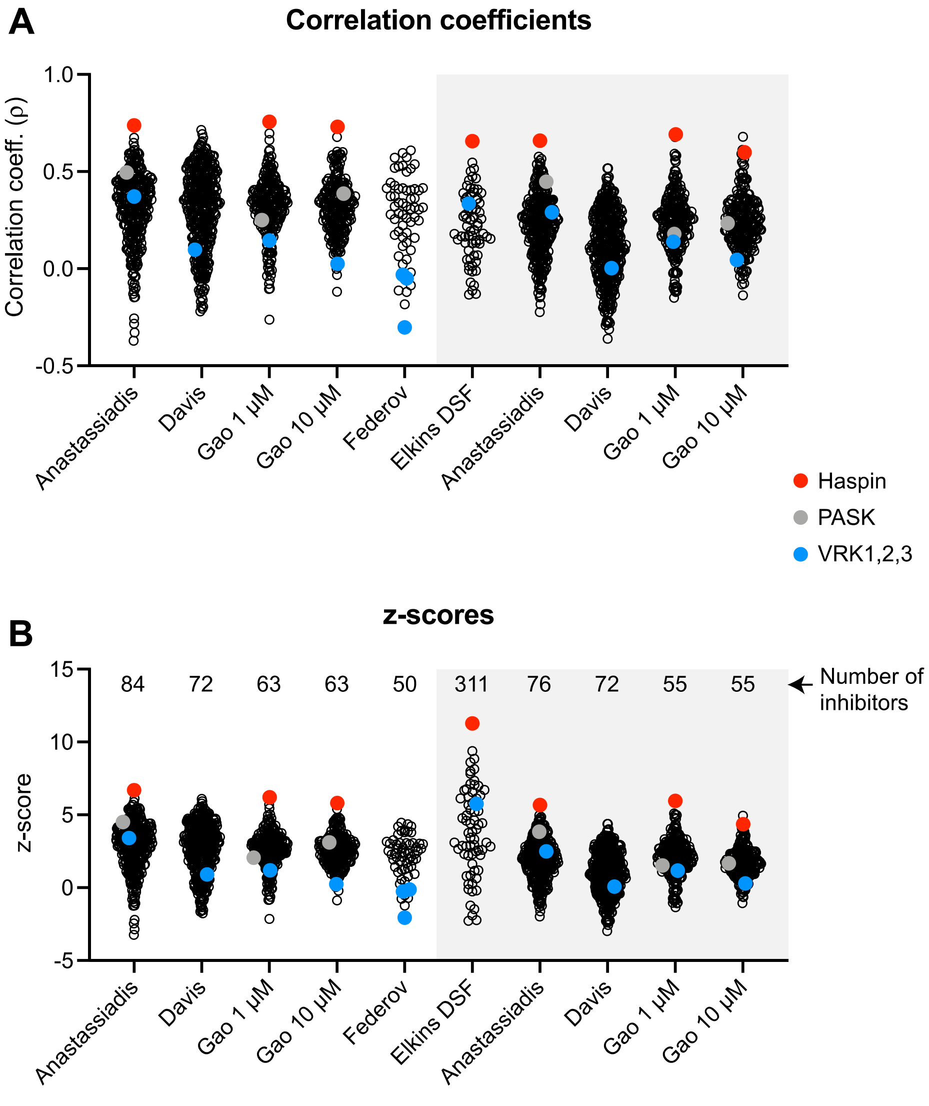


**Figure S1**

Results of KiPIK screens for mitotic H3T3 kinases in HeLa cell extracts.

**A**. Correlation coefficients for all kinases tested. **B**. z-scores for all kinases tested.

The screens shown on a gray background were previously published (48). Z-scores allow a more direct comparison of the robustness of kinase ranking between screens which accounts for the number of inhibitors used (shown at the top of the plot), as described (48).

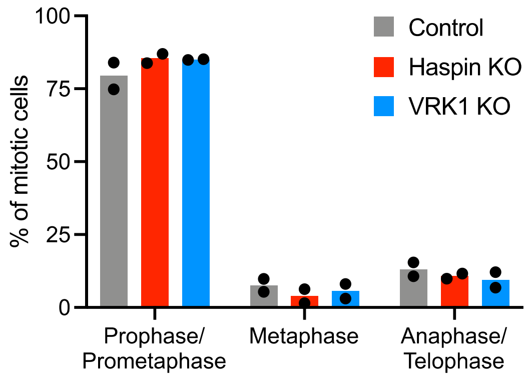


**Figure S2**
Control, Haspin KO, and VRK1 siRNA-treated HeLa cells were fixed and stained with DAPI to visualize DNA, and with antibodies to H3T3ph and H3S10ph (n = 2 independent experiments). The stage of mitosis was determined by DAPI staining for all cells from prometaphase to early telophase.

### A GSG2 (Haspin) gene (2 bp insertion in exon 1)

```
       100                                     110                                     120
        R  W  K  L  R  A  R  P  S  L  T  V  T  P  R  R  L  G  L  R  A  R  P  P  Q     K  C
WT    AGCGCTGGAAGCTGCGAGCTCGCCCAAGCCTAACCGTGACCCCAAGACGCCTGGGGCTGCGAGCTCGGCCCCCGC--AGAAGTGC
      |||||||||||||||||||||||||||||||||||||||||||||||||||||||||||||||||||||||||||  ||||||||
KO    AGCGCTGGAAGCTGCGAGCTCGCCCAAGCCTAACCGTGACCCCAAGACGCCTGGGGCTGCGAGCTCGGCCCCCGCAAAGAAGTGC
        R  W  K  L  R  A  R  P  S  L  T  V  T  P  R  R  L  G  L  R  A  R  P  P  Q  R  S  A

                130                                     140                                     150
      S  T  P  C  G  P  L  R  L  P  P  F  P  S  R  D  S  G  R  L  S  P  D  L  S  V  C  G  Q
WT    AGCACACCCTGCGGCCCGCTCCGACTTCCGCCCTTCCCCAGCCGCGACTCCGGCCGCCTCAGCCCGGACCTCAGCGTGTGCGGCC
      |||||||||||||||||||||||||||||||||||||||||||||||||||||||||||||||||||||||||||||||||||||
KO    AGCACACCCTGCGGCCCGCTCCGACTTCCGCCCTTCCCCAGCCGCGACTCCGGCCGCCTCAGCCCGGACCTCAGCGTGTGCGGCC
       A  H  P  A  A  R  S  D  F  R  P  S  P  A  A  T  P  A  A  S  A  R  T  S  A  C  A  A

                       160                                     170                                     180
        P  R  D  G  D  E  L  G  I  S  A  S  L  F  S  S  L  A  S  P  C  P  G  S  P  T  P  R
WT    AGCCCAGGGACGGCGACGAGCTGGGCATCAGTGCCTCCCTGTTCAGCTCTCTGGCCTCGCCCTGCCCCGGGTCCCCAACGCCAAG
      |||||||||||||||||||||||||||||||||||||||||||||||||||||||||||||||||||||||||||||||||||||
KO    AGCCCAGGGACGGCGACGAGCTGGGCATCAGTGCCTCCCTGTTCAGCTCTCTGGCCTCGCCCTGCCCCGGGTCCCCAACGCCAAG
      S  P  G  T  A  T  S  W  A  S  V  P  P  C  S  A  L  W  P  R  P  A  P  G  P  Q  R  Q  G

                            190                                     200                                     210
       D  S  V  I  S  I  G  T  S  A  C  L  V  A  A  S  A  V  P  S  D  L  H  L  P  E  V  S
WT    GGACAGTGTCATCTCGATCGGCACCTCCGCCTGTCTGGTTGCAGCCTCAGCCGTCCCGAGCGGCCTCCACCTCCCAGAAGTCTCC
      |||||||||||||||||||||||||||||||||||||||||||||||||||||||||||||||||||||||||||||||||||||
KO    GGACAGTGTCATCTCGATCGGCACCTCCGCCTGTCTGGTTGCAGCCTCAGCCGTCCCGAGCGGCCTCCACCTCCCAGAAGTCTCC
        T  V  S  S  R  S  A  P  P  P  V  W  L  Q  P  Q  P  S  R  A  A  S  T  S  Q  K  S  P

                                 220                                     230                                     240
      L  D  R  A  S  L  P  C  S  Q  E  E  A  T  G  G  A  K  D  T  R  M  V  H  Q  T  R  A  S
WT    CTGGACCGAGCATCTCTCCCCTGCTCCCAGGAGGAAGCGACAGGAGGAGCCAAGGACACCAGGATGGTCCACCAAACCCGCGCCA
      |||||||||||||||||||||||||||||||||||||||||||||||||||||||||||||||||||||||||||||||||||||
KO    CTGGACCGAGCATCTCtcccctgctcccaggaggaagcgacaggaggagccaaggacaccaggatggtccaccaaacccgcgcca
       W  T  E  H  L  S  P  A  P  R  R  K  R  Q  E  E  P  R  T  P  G  W  S  T  K  P  A  P

                                      250
        L  R  S  V  L  F  G  L  M  N  S  G  T  P  E  D  S
WT    GCCTCAGGTCAGTTCTCTTTGGCCTTATGAACTCAGGAACCCCTGAGGATT
      |||||||||||||||||||||||||||||||||||||||||||||||||||
KO    gcctcaggtcagttctctttggccttatgaactcaggaacccctgaggatt
      A  S  G  Q  F  S  L  A  L  *
```

### B VRK1 gene (11 bp deletion in exon 5)

```
                                                   100                                     110
                                Q  K  W  I  R  T  R  K  L  K  Y  L  G  V  P  K  Y  W  G  S
WT    ACAAAGATCTGTTTTAATTTGTAGTTCAGAAATGGATTCGTACCCGTAAGCTGAAGTACCTGGGTGTTCCTAAGTATTGGGGGTC
      ||||||||||||||||||||||||||||||||||||||||||||||||||||           ||||||||||||||||||||||
KO    ACAAAGATCTGTTTTAATTTGTAGTTCAGAAATGGATTCGTACCCGTAAGCT-----------GTGTTCCTAAGTATTGGGGGTC
                                Q  K  W  I  R  T  R  K  L                 C  S  *

                120
      G  L  H  D  K  N  G  K  S
WT    TGGTCTACATGACAAAAATGGAAAAAGGTAAAAATATGT
      |||||||||||||||||||||||||||||||||||||||
KO    TGGTCTACATGACAAAAATGGAAAAAGGTAAAAATATGT
```

**Figure S3**

DNA and predicted amino acid sequences of Haspin and VRK1 genes in wild type (WT C631) HAP1 cells (top lines) compared with **A**, Haspin KO (HZGHC000047c016) and **B**, VRK1 KO (HZGHC000073c014) HAP1 cells (bottom lines). Numbering refers to amino acid position. Grey shaded regions are exons. Red regions highlight deletions or insertions resulting in frame shifts. Sequencing data from Horizon Discovery.

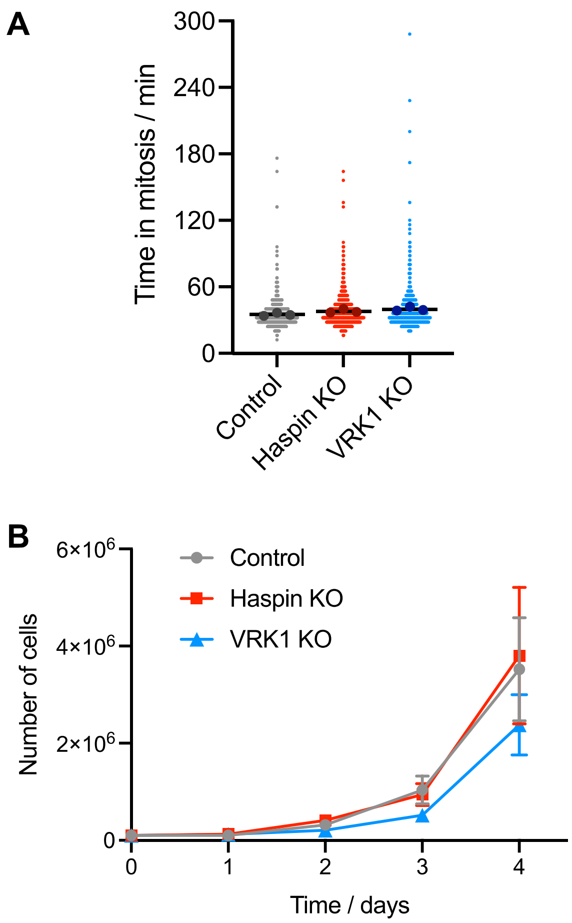


**Figure S4**

In **A**, the duration of mitosis (from nuclear envelope breakdown to anaphase onset) in control, Haspin KO and VRK1 KO HAP1 cells was determined by live imaging. Between 145 and 342 cells per condition were evaluated from three separate fields of view in each of three independent experiments. Small symbols represent individual cells, and large symbols show the mean duration of mitosis in each of the three experiments. Black bars show the means of these means (n = 3 independent experiments). In **B**, the proliferation of control, Haspin KO and VRK1 KO HAP1 cells was measured by cell counting (n = 3 independent experiments).

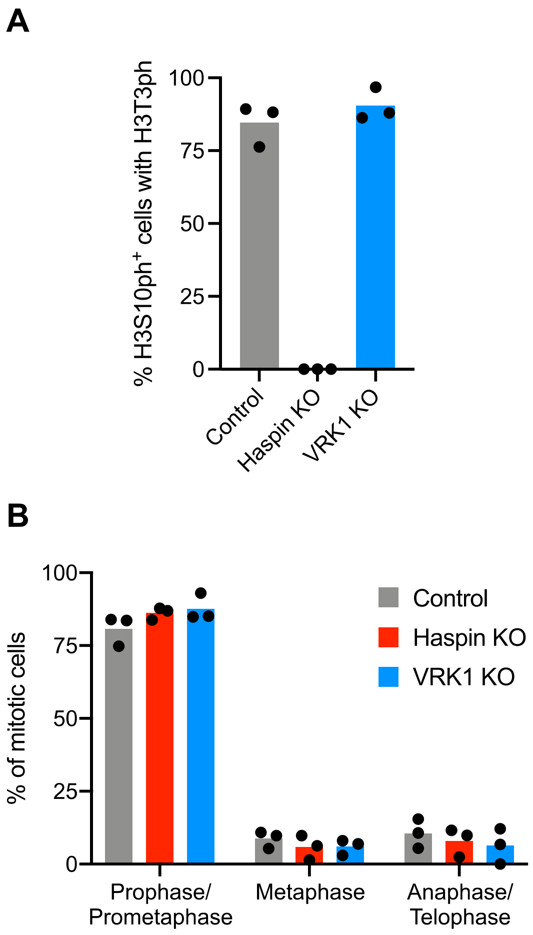


**Figure S5**

In **A**, control, Haspin KO and VRK1 KO HAP1 cells were fixed and stained with DAPI to visualize DNA, and with antibodies to H3T3ph and H3S10ph. The percentage of all cells with H3S10ph staining that had detectable H3T3ph was scored (n = 3 independent experiments). In **B**, the stage of mitosis was determined by DAPI staining for all cells from prometaphase to early telophase (n = 3 independent experiments).
